# Transcriptomic profiling of unmethylated full mutation carriers implicates TET3 in FMR1 CGG repeat expansion methylation dynamics in Fragile X syndrome

**DOI:** 10.1101/2024.10.21.617801

**Authors:** Grace Farmiloe, Veronika Bejczy, Elisabetta Tabolacci, Rob Willemsen, Frank Jacobs

**Author notes:** Corresponding author, Frank Jacobs.

## Abstract

**Background:** Fragile X syndrome (FXS) is a neurodevelopmental disorder caused by the expansion of a CGG repeat in the 5’UTR of the FMR1 (fragile X messenger ribonucleoprotein 1) gene. Healthy individuals possess a repeat 30-55 CGG units in length. Once the CGG repeat exceeds 200 copies it triggers methylation at the locus. This methylation covers the FMR1 promoter region and silences expression of the gene and the production of FMRP (fragile X messenger ribonucleoprotein). The loss of FMRP is responsible for a number of pathologies including neurodevelopmental delay and autism spectrum disorder. Methylation of the expanded repeat in the FMR1 locus is the causal factor for FXS, however it is not known why the expanded repeat triggers this epigenetic change or how exactly DNA methylation is established. Intriguingly, genetic engineering of expanded CGG repeats of over 300x in the FMR1 locus in mice remains unmethylated. Also in humans, in very rare cases, individuals can have an FMR1 CGG expansion >200x but the locus remains unmethylated. These unmethylated full mutation individuals give us a rare opportunity to investigate the mechanism of FMR1 promoter methylation.

**Methods:** Fibroblasts were obtained from a healthy control, an FXS patient and two unmethylated full expansion carriers. RNA was extracted and comparative transcriptomic analysis was performed on all samples. Whole genome sequencing was carried out on DNA from the two UFM carriers and the results analysed to investigate DNA variants that could explain the observed differences in gene expression.

**Results:** Our analyses focused on genes involved in epigenetic modification. We show that Tet methylcytosine dioxygenase 3 (TET3), a gene involved in DNA methylation, is significantly downregulated in UFM carriers compared to healthy controls or FXS patient derived cells. Genomic analyses reveal a number of rare variants present in the TET3 locus in UFM carriers when compared to the reference genome. No single variant has a significant predicted effect, raising the possibility that a trans acting variant could be driving the differential gene expression.

**Conclusion:** Our results suggest that TET3 is a candidate factor responsible for the lack of methylation of the expanded FMR1 locus. Further analyses are needed to further elucidate this relationship, however given its potential to directly interact with CGG repeats and its ambiguous role in 5-hydroxy-methylation of CG containing sequences, TET3 is a strong candidate for further exploration.

## Background

Fragile X syndrome (FXS) is a neurodevelopmental disorder caused by the expansion of a CGG simple repeat in the 5’ untranslated region (5’UTR) of the fragile X messenger ribonucleoprotein 1 (FMR1) gene. In the healthy population, repeat numbers range between 30-55x CGG repeats. Individuals carrying 56-200x repeats are said to carry a pre-mutation (PM) (1). In some but not all cases of PM, the lengthened CGG repeat can cause degenerative disorders such as Fragile X-associated tremor/ataxia syndrome (FXTAS) or fragile X associated premature ovarian insufficiency (FXPOI) (2–6). In individuals affected by fragile X syndrome, repeat lengths exceed a threshold of 200x CGG repeats (7). Once the repeat exceeds 200x it gains DNA methylation which shuts down expression of FMR1 and leads to a global loss of fragile X messenger ribonucleoprotein (FMRP), an RNA-binding protein which plays an essential role in brain development (8). Fragile X individuals who lack FMRP exhibit symptoms including neurodevelopmental delay and autism spectrum disorder. Interestingly, at very early stages of development, there is some evidence that the FMR1 gene in FXS individuals is still active (9). The locus is thought to gain methylation between 6-11 weeks of embryonic development (10). Indeed, FXS embryonic stem cells (ESCs) carrying a full mutation which has not yet become methylated have been shown to gain methylation at the locus after 60 days of neuronal differentiation (11). This shows that methylation at the expanded repeat is dynamic and suggests that another factor, in addition to the repeat expansion, is responsible for establishing DNA methylation at the FMR1 locus.

Humanised FXS mice, which have an engineered CGG repeat expansion in the FMR1 locus were created to study FXS and the methylation dynamics at the locus. Surprisingly, these mice do not gain methylation at the FMR1 locus, even with repeat lengths up to 341 (12–15). These mice display the pre-mutation associated symptoms but escape full methylation. This discrepancy with human FXS patients further supports the hypothesis that the repeat expansion by itself is insufficient to explain the silencing of the FMR1 gene. This raises the possibility that mice lack a certain cellular factor required for establishing DNA methylation at the FMR1 locus.

Further evidence for the presence of a cellular factor implicated in the methylation at the FMR1 expanded repeat comes from a number of rare individuals who possess a full CGG expansion mutation but never gain methylation at the locus (16–18). These unmethylated full mutation (UFM) individuals are very rare, with only a handful discovered globally. It would seem that these individuals lack the elusive factor responsible for establishing methylation at the expanded CGG repeat, without causing noticeable differences in the individual’s phenotype (16–18). Interestingly, in these individuals there is also evidence of a dynamic methylation landscape at the FMR1 locus. In some cases, during the creation of induced pluripotent stem cells (iPSCs), cells from these individuals can gain methylation at the locus (19,20). This has been linked to instability in the CGG repeat itself; in one instance the gaining of methylation was only seen in clones which also experienced further expansion of the CGG repeat (19). Both UFM and PM carriers have higher FMR1 expression levels than those within the 30-55x repeat range but counter-intuitively, lower FMRP levels (21). This molecular phenotype sometimes does not affect the individual however in some cases it can lead to FXTAS or FXPOI in PM individuals (5,22,23).

These findings suggest that like the humanised FXS mice, perhaps human UFM carriers lack a particular factor required for establishing DNA methylation at the FMR1 locus. In this study, we profile the transcriptomes of two unrelated UFM carriers, a full mutation (FM) carrier and a healthy control to investigate this phenomenon. We focus on genes with epigenetic modifying function and particularly those involved in DNA methylation to see if there is any differential expression that could explain the lack of methylation in UFM carriers. The results of our analyses reveal TET3 as a promising candidate. It has DNA binding capacity, it plays an important role in DNA methylation and is significantly downregulated in UFM carriers.

## Methods

### Cell culture of fibroblasts

4 fibroblast cell lines were cultured for this study, 1 healthy control with a repeat length <55x, 1 FXS control with a methylated CGG repeat >200x and 2 UFM carriers with unmethylated CGG repeats >200x. Each fibroblast line was cultured in triplicate to create three experimental replicates for RNA extraction and sequencing. One culture well of each UFM cell line was harvested for DNA extraction and whole genome sequencing Fibroblasts were cultured in DMEM + Glutamax (Gibco) with 15% HIFBS (Gibco), 0.1 M non-essential amino acids (Gibco), 100 U/mL penicilin-streptomicin (pen/strep, Gibco™). Cells were kept in the incubator at 37°C and 5% CO2. Medium was changed every other day and cells were passed once a week using 0.25% trypsin (Gibco) + 0.01% EDTA.

### RNA extraction

RNA from fibroblasts was isolated in 400 µl TRIzol Reagent (Invitrogen™) according to manufacturer’s recommendations. Samples were treated with DNAseI (Roche) and cleaned using the DNA Clean & Concentrator™-5 Kit (ZYMO Research). Libraries for the UFM 1 samples were prepared with the TruSeq Stranded Total RNA (Illumina) with Ribo-Zero ribosomal RNA depletion, and sequenced paired-end, 75bp on a NextSeq 550 system (Illumina) by MAD: Dutch Genomics Service & Support Provider (Swammerdam Institute for Life Sciences, Amsterdam). The UFM 2 RNA samples were sent for library prep and sequencing at genomescan (https://www.genomescan.nl/).

### RNA-seq data analysis

RNA data was analysed using the public Freiburg Galaxy server (24,25), usegalaxy.eu). Trimmomatic was used to remove adapters and trim reads (26) version 0.36.5 for paired-end reads (ILLUMINACLIP TruSeq3 paired-end), the cut option was set if average per base quality in a 4-base sliding window was below 20, and reads below 30 bases were dropped. Reads were mapped to the built-in reference genome hg38 using HISAT2 (27) (Galaxy Version 2.1.0+galaxy3). Reads were assigned to gencode V36 features using featureCounts (28) (Galaxy Version 1.6.3) with -p, -d 75 -D 900 -B -C. The featureCounts output was analysed using DESeq2 (29) (Galaxy Version 2.11.40.3) with default settings. De novo transcript assembly was performed using stringtie (Galaxy Version 2.1.1) without using a reference guide (30). bamCoverage was used to generate coverage tracks for both strands (Galaxy Version 3.0.2.0, with deepTools2 (Version 3.0.2) and samtools (Version 1.7)) from the deeptools2 package (31) (bin size 1). Coverage tracks were scaled on UCSC with a scaling factor based on the number of uniquely assigned reads from HISAT2. Replicates were then gathered into track collections on the UCSC and scaled to the replicate with the highest number of uniquely assigned reads from featureCounts, these track collections were used for visualisation. Volcano plots were made in R using the ggplot2 package (32).

### DNA extraction and whole genome sequencing

DNA was extracted using Quick-DNA miniprep plus kit™ (ZYMO Research) according to the manufacturer instructions. DNA was then sent to the Hartwig institute for library preparation and sequencing using the NovaSeq6000 system (Illumina).

### Whole genome sequencing data analysis

Whole genome sequencing data was analysed using the public Freiburg Galaxy server ((24,25), usegalaxy.eu). Adapters were removed and reads were trimmed using Trimmomatic (26) version 0.36.5 for paired-end reads (ILLUMINACLIP TruSeq3 paired-end), the cut option was set if average per base quality in a 4-base sliding window was below 20, and reads below 30 bases were dropped. Reads were then mapped against built in genome hg38 using BWA MEM (Galaxy version 0.7.17.1) (33) using default settings. The resulting BAM file was sorted by coordinate using SortSam (Galaxy version 2.18.2.1) from the SAMtools package (34) and default settings. Variant calling was performed using FreeBayes (Galaxy version 1.3.1) (35) against the built in hg38 reference genome. The resulting VCF file was then filtered using SNPSift filter from the SNPeff package (Galaxy version 3.4) (36), variants with a read depth below 10 and a quality score below 30 were excluded. The VCF file was then normalised using bcftools norm (Galaxy version 1.10) (37) and annotated with the dbSNP variant names using SNPSift annotate (Galaxy version 4.3+t.galaxy1) (36). The VCF file was then visualised on the UCSC genome browser (38). Ensembl’s Variant Effect Predictor and the dbSNP database were also used to help annotate variants (39,40).

### Expression analysis in organoid data from human and rhesus

For the comparison of Human and rhesus cortical organoid development, published data from Field et al. (41) was used (GEO: GSE106245) (41). Data was accessed in a summarised form and basemean values, which had been generated from 2 replicates, were plotted on the graph.

### Generation of FXS organoids

FXS iPSCs were grown in StemFlex medium (Gibco). Medium was changed every 2-3 days and cells were passaged every 3-4 days using Versene (ThermoFisher) and medium supplemented with Y-27623 ROCK inhibitor (Tocris). Cells were kept in the incubator at 37°C and 5% CO_2_. The organoid differentiation protocol was based on the methods of Eiraku et al. 2008 (42). To make embryoid bodies Aggrewell plates were used (Stemcell technologies). 6 million iPSCs were seeded per well in medium supplemented with Y-27623. 24hrs after seeding cells were transferred to an ultra low attachment 60mm plate (Corning) in neuronal differentiation medium (DMEM-F12 (Gibco) supplemented with 20% KnockOut Serum Replacement (Gibco), 100 U/ml penicillin/100 µg/ml streptomycin (Gibco), 2 mM GlutaMAX (Gibco), 1x MEM Non-Essential Amino Acids solution (Gibco), 100 µM 2-mercaptoethanol (Gibco)) with 8 ng/ml fresh bFGF (Sigma) and 1 mM sodium pyruvate (Gibco)). 24 hours after transfer medium was refreshed with differentiation medium with freshly added 3 µM IWR-1-Endo, 1 µM Dorsomorphin, 10 µM SB-431542 hydrate, and 1 µM Cyclopamine hydrate (day 0 of differentiation), Medium was refreshed with fresh inhibitors every other day. On day 3 of differentiation, organoids were moved to a shaker to prevent clumping. On day 18, the medium was changed to Neurobasal/N2 medium (Neurobasal (Gibco) supplemented with 100 U/ml penicillin/100 µg/ml streptomycin (Gibco), 2 mM GlutaMAX (Gibco), 1x N-2 supplement (Gibco)) supplemented with 1 µM Cyclopamine hydrate. From D24, no inhibitors were added anymore to the medium until organoids were harvested. Organoids were grown for 60 days and harvested at day 12, day 36 and day 60.

### RNA extraction and qPCR

RNA was isolated using TRIzol (Invitrogen™) following the manufacturer’s instructions. Potential DNA contamination was removed with DNAseI (Roche) and samples were cleaned using the DNA Clean & Concentrator™-5 Kit (ZYMO Research). qRT-PCR was performed using the QuantiTect SYBR Green RT-PCR Kit (Qiagen) with 3 technical replicates for each biological replicate sample in 10ul reactions. The following primers were used, GAPDH was used as a housekeeping gene: TET3_qRT-PCR_F1: CCCACGGTCGCCTCTATC, TET3_qRT-PCR_R1: CTCCTTCCCCGTGTAGATGA, GAPDH_qRT_PCR_01_F1: AATCCCATCACCATCTTCCA, GAPDH_qRT_PCR_01_R1: TGGACTCCACGACGTACTCA

### Multiple sequence alignment of TET3

TET3 isoform protein sequences for human, monkey and mouse were downloaded from UCSC genome browser (38). Protein sequences from annotated transcripts (ENST00000409262.8, ENST00000305799.8, ENSMUST00000089622.11, ENSMUST00000186548.7, ENSMUST00000190295.2, ENSMMUT00000028916.3, ENSMMUT00000028918.4, ENSMMUT00000075204.2) were aligned using clustal omega and default settings (43). The alignment was then manually annotated based on information published in Iyer et al., 2009 (44).

### Comparison of human and mouse expression during brain development

Human brain developmental expression data was downloaded from the Brainspan atlas of the developing human brain (https://www.brainspan.org/) for both FMR1 and TET3. Mouse brain developmental expression data for FMR1 and TET3 was downloaded from the EBI expression atlas, specifically experiment ‘E-MTAB-6798: Mouse RNA-seq time-series of the development of seven major organs’ (https://www.ebi.ac.uk/biostudies/arrayexpress/studies/E-MTAB-6798). The mouse data points were then adjusted to match human developmental time points using the ‘translating time’ website, https://www.translatingtime.org/ (45). Due to the differences in scale across all of the data points from each dataset, the average TPM values across human and mouse were calculated and used to scale the data. These data points were then plotted on a dual axis graph to display the expression pattern over time for the two species.

## Results

### Inter-individual and inter-species differences in methylation dynamics of FMR1 CGG repeat expansions

The length of the FMR1 CGG repeat is a continuous variable and individuals with repeat lengths that fall between different categories based on repeat size have very different molecular phenotypes. There are 4 categories of repeat size and relative FMRP expression levels: The majority of individuals have approximately 30 CGG repeats and normal levels of FMRP expression, PM carriers have 56-200 CGG repeats and lowered FMRP expression, FXS individuals have >200 repeats and no FMRP expression and, very rarely, UFM carriers have >200 repeats and expression levels similar to those found in PM carriers (**Figure 1**). It is interesting to note that CGG repeat lengths which fall around the boundaries of these categories can be variable as to which molecular phenotype they present. Interestingly, mouse models with an engineered CGG expansion in FMR1, even to the length of 341 repeats, do not gain methylation and share a molecular phenotype with PM and UFM carriers (**Figure 1**, (12–15)). This differential methylation characteristic between species implies that the driver behind methylation in humans at the locus is a gene or factor involved in DNA methylation or other epigenetic modifications. Given the differences between methylation between mouse and human we would also expect there to be differences in a candidate gene between these species. Based on the above, we generated a candidate gene list of 44 DNA and histone modifiers (**Supplemental table 1**).

**Figure 1:**
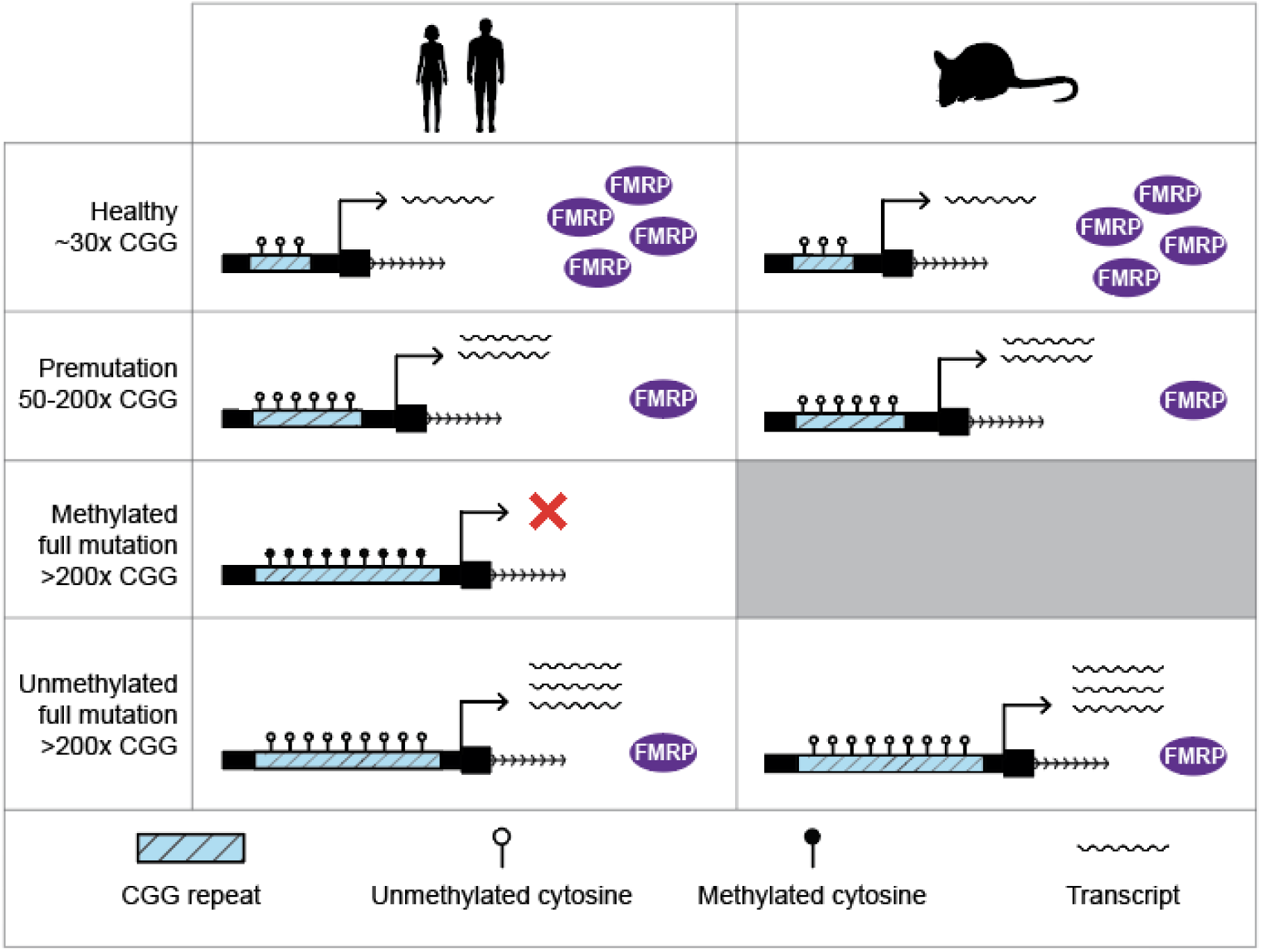
Variation in the FMR1 promoter and expression in human and mouse. Observed variation in CGG repeat length, methylation status, transcript and protein levels in human (on the left) and mouse (on the right).

### Transcriptomic profiling confirms the UFM molecular phenotype at the *FMR1* locus

The lack of methylation in the FXS mouse model suggests that there is a difference in the regulation of methylation between mouse and human at the FMR1 locus. The existence of rare UFM carriers offers an unparalleled opportunity to investigate the factors behind the methylation of the CGG repeat expansion unencumbered by the issues associated with inter-species comparison. UFM carriers are quite rare in the human population, but we were able to obtain two fibroblast cell lines generated from genetically unrelated UFM carriers which allowed us to perform a comparative transcriptomic analysis of UFM carriers, FXS patients and healthy controls. The first UFM carrier cell line (UFM 1), contains an unmethylated CGG repeat of approximately 233 repeats in the FMR1 5’UTR (16,20). As most expansion carriers possess a mosaic of repeat lengths, this fibroblast cell line has undergone selection to ensure a narrow range of repeat lengths around 230x. The second UFM carrier cell line (UFM 2) has also been previously reported (17–19) and contains a range of 265–430 repeats.

Transcription at the FMR1 locus has previously been studied in cells from individuals with different repeat lengths using qPCR (16,18–20). However, a full transcriptional profile of the locus in UFM carriers has not previously been generated. The transcriptional profile at the FMR1 locus confirms the previously reported molecular phenotypes for the 3 types of cell lines sequenced: Compared to healthy control fibroblasts, FMR1 transcript levels were upregulated in both UFM lines and FMR1 transcripts were completely absent in the FXS cell line. (**Figure 2b, d**). The RNA-seq analysis also revealed high levels of a transcript originating from the FMR1 5’UTR and continuing into the first intron in the UFM carriers. This transcript encodes only exon 1 and intron 1 of the FMR1 gene, and lacks a clear protein-coding potential, when translated the resulting potential protein has a stop codon after 50 amino acids. *De novo* transcriptome assembly and annotation reveals that this transcript is expressed from the CGG-repeat, and is also observed at very low levels in healthy control fibroblasts (**Figure 2a, b**) which is further supported by publicly available RNA-seq expression data (**Supplemental figure 1**). Low levels of these transcripts have been identified in PM carriers and the translation of these aberrant transcripts has been linked to FXTAS (46). Interestingly, low levels of the expanded CGG driven transcript have also occasionally been identified in FXS individuals and have been linked to the triggering of methylation at the locus (11). This further demonstrates the complexity of the methylation dynamics at the FMR1 locus; the presence of high levels of the CGG-driven transcripts in UFM carriers implies that it may be necessary but not sufficient to trigger methylation.

**Figure 2:**
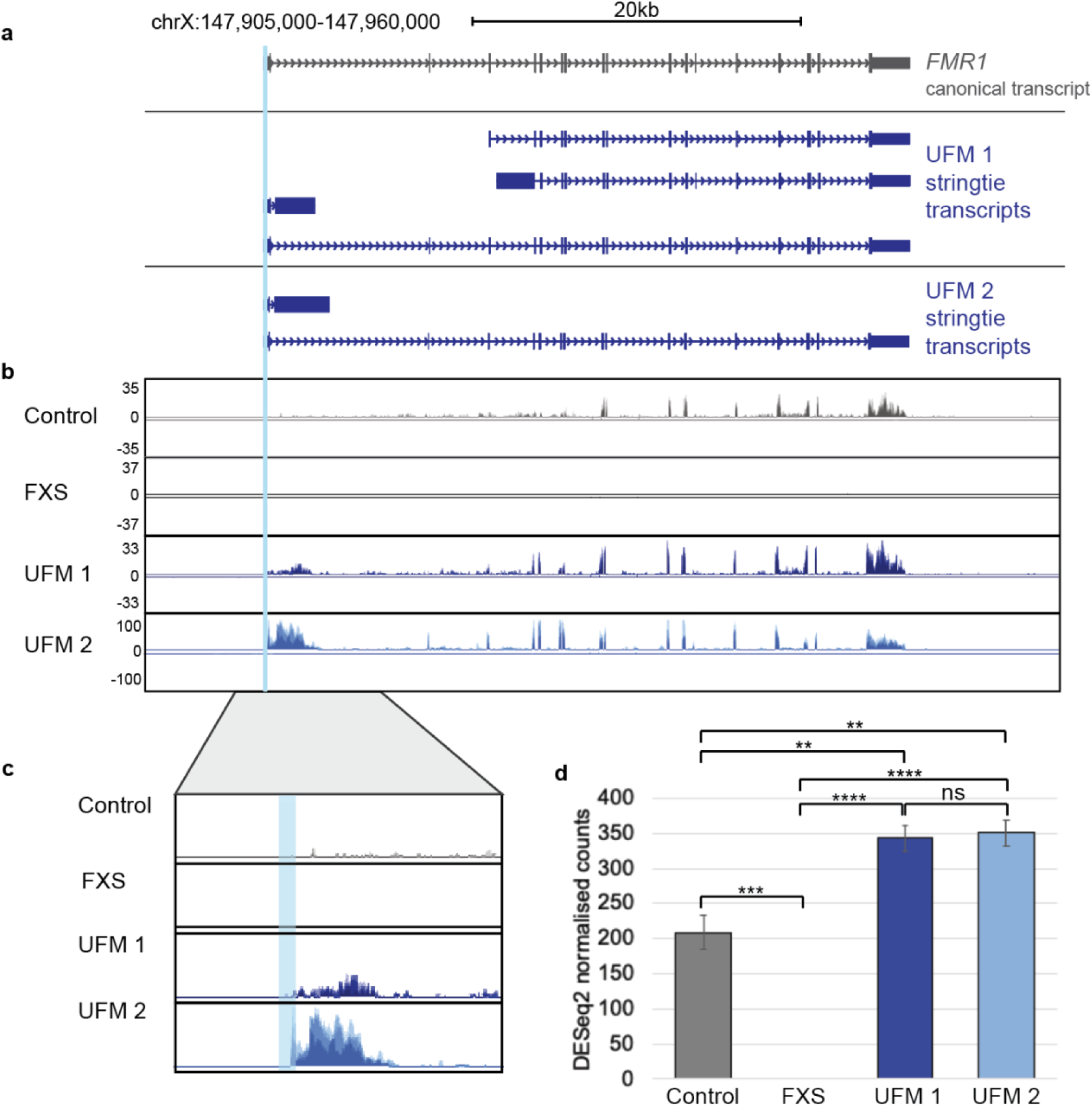
FMR1 expression in an unmethylated expansion carrier. (a) Schematic of the FMR1 locus, the canonical transcript in grey and stringtie de novo assembled transcripts generated using RNA-seq data from fibroblasts from an FMR1 UFM carrier in blue. (b) Coverage plots of RNA-seq data at the FMR1 gene from fibroblasts obtained from a control individual, an FXS patient and two UFM carriers. Coverage tracks are scaled based on the total number of reads successfully mapped to the genome. Coverage tracks were generated from track collections and show merged tracks from 3 replicates. (c) Detailed comparison of the control and UFM RNA-seq data at intron 1. Showing below a representation of the transcript and potential translation (d) Barplot showing DESeq2 normalised read counts for FMR1 for the control, FXS and UFM cell lines; 1-way ANOVA, Tukey’s multiple comparison test: **** P < .0001, *** P<0.001, ** P< .01.

### Analysis of epigenetic modifiers reveals TET3 as a candidate gene

We next investigated differences in the expression of candidate epigenetic modifying genes between the two UFM carriers, a healthy and an FXS control cell line. For the analysis, the three replicates for UFM 1 and the three replicates for UFM 2 were grouped and compared in turn to the healthy control samples (**Figure 3a**) and the FXS samples (**Figure 3b**). After the filtering criteria of an adjusted p-value <0.01 and a fold change greater than 1.5, 35 out of the 44 candidate genes were expressed at high enough levels to compare with the control and FXS samples. The normalised reads for the top three genes from each comparison based on p-value and fold change from the DESeq2 analysis were further investigated (**Figure 3c**). When compared against the healthy control both TET3 and TET1 were downregulated with a fold change greater than 1.5x and an adjusted p-value <0.01 in the UFM carriers and SET Domain Containing 1B (SETD1B) was upregulated. In comparison with the FXS individual TET3 and Enhancer Of Zeste 2 Polycomb Repressive Complex 2 Subunit (EZH2) passed the thresholds and were downregulated in the UFM carriers.

**Figure 3:**
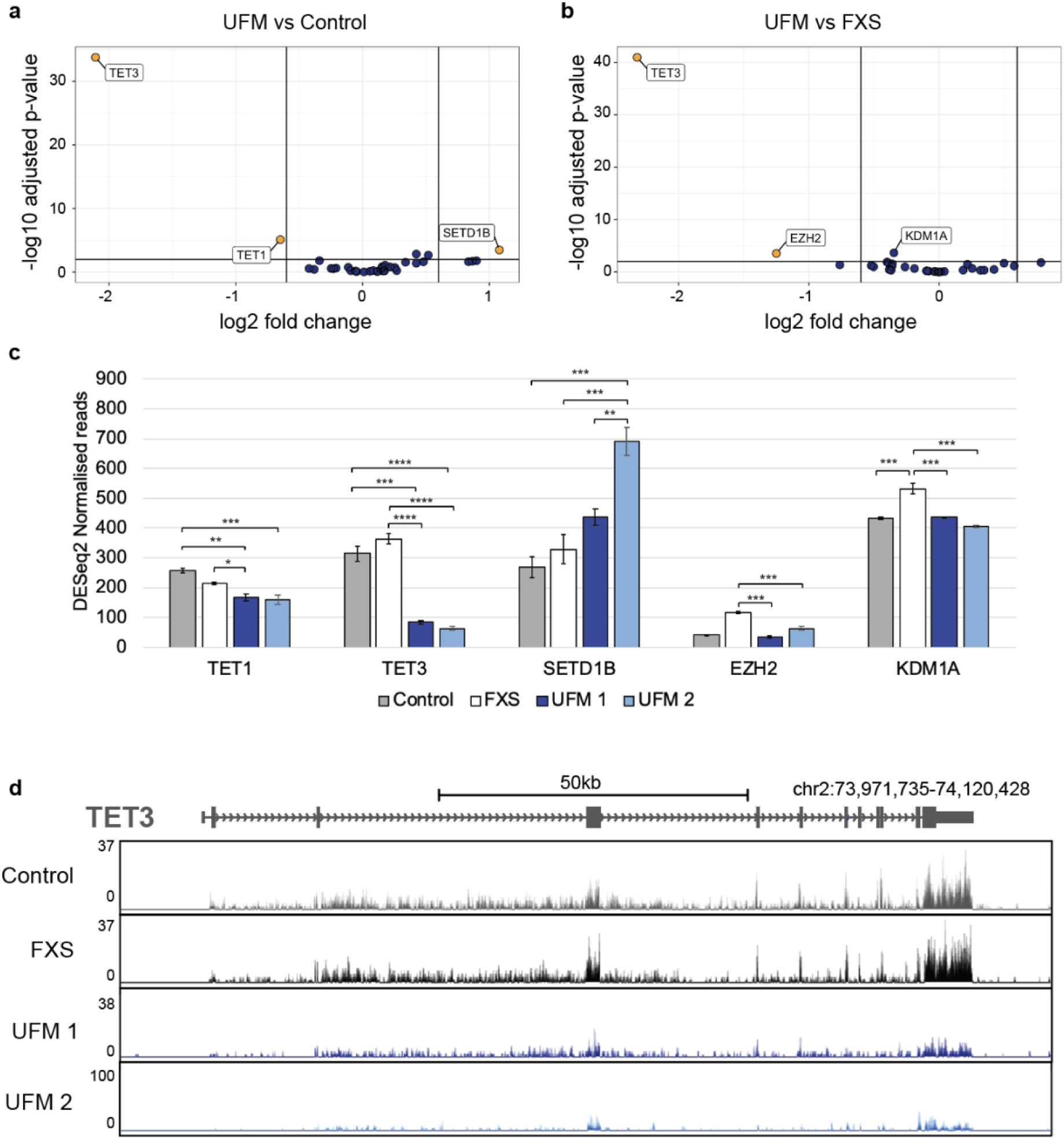
Candidate gene expression in unmethylated expansion carriers. (a, b) Volcano plots showing Log2 fold change and significance from DESeq2 analysis of candidate genes comparing the gene expression of both UFM lines with the FXS and healthy control lines. Orange points represent genes that pass the threshold of fold change >1.5 and p-value <0.01, blue points did not pass this threshold. (c) Barplots showing DESeq2 normalised read counts for differentially expressed candidate epigenetic modifiers for the control, FXS and UFM cell lines; 1-way ANOVA, Tukey’s multiple comparison test: **** P < 0.0001, *** P <0.001, ** P < .0.01, * P < 0.05, ns - not significant. (d) Mapped RNA-seq data showing the raw read data for TET3.

Upon comparison with both the FXS and unexpanded control we see a robust downregulation of ~80% of TET3 in both UFM carriers. This strong downregulation can also clearly be seen in the mapped reads from the RNA-Seq, (**Figure 3d**) with very low levels of TET3 expression in the UFM lines when compared to control and FXS lines. RNA-seq also reveals that differential expression in UFM carriers is due to an overall decrease of TET3 expression, and not the loss of any particular TET3 transcript isoform

To rule out the possibility that the differential gene expression we observed between the two UFM carriers, the control and the FXS lines is due to the coincidental aberrant expression of these genes in healthy controls and FXS cells, we confirmed in two, unrelated, healthy control fibroblast lines that expression levels of TET3 were similar to the level of the healthy and FXS control cells we used for the initial comparisons (**Supplemental figure 2**). Overall, our data points towards a possible involvement of TET3 in the resilience against DNA methylation displayed in both UFM 1 and UFM 2. Because of the ability of TET3 to bind directly to CGG repeat, its described role in DNA methylation and the aberrant expression observed in UFM carrier fibroblasts, TET3 is a candidate contributor to the pathological CGG methylation of the FMR1 promoter in FXS patient cells.

### WGS reveals no potential cis acting variants to influence TET3 differential expression

It is possible that the observed differences in expression of TET3 in the UFM cell lines are caused by structural variation or a cis acting genetic variant in the coding or non-coding regions of the TET3 locus. To assess whether this is the case, we performed high coverage (~30x) whole genome sequencing for both UFM cell lines. When compared to the reference genome, both UFM genomes showed a large number of SNPs and small indels. Variant calling was performed for the TET3 locus covering approximately 600kb for both UFM I and UFM II genomes (**Figure 4a**). We hypothesised that the variant influencing TET3 expression levels would be rare (MAF <0.01) and shared between the two UFM carriers. The variants were filtered for those that were present in both samples and the allele frequencies for the variants were retrieved from gnomAD (https://gnomad.broadinstitute.org/) and dbSNP (https://www.ncbi.nlm.nih.gov/snp/) (40,47).

**Figure 4:**
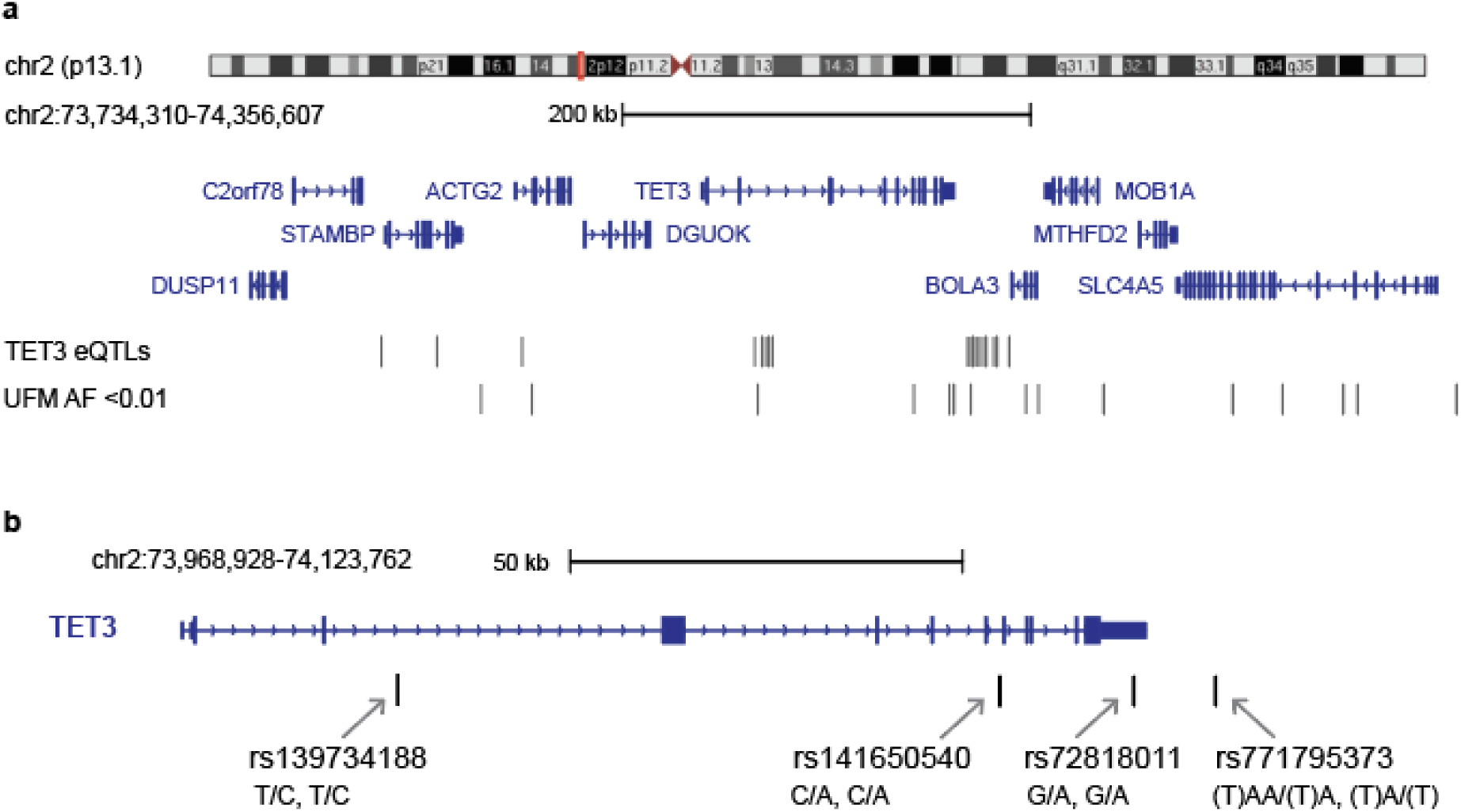
TET3 variants and expression in the general population. a. Genomic variants for TET3 for both UFM cell lines in the TET3 locus ~600kb showing the genes, TET3 eQTLs and SNPs found in both UFM carriers with a MAF <0.01 and mapped RNA-sequencing coverage in each line at the locus. b. The 4 SNPs in or adjacent to TET3, showing 2 in the introns, one in the 3’UTR and one just downstream. The Genotypes for UFM1/UFM2 are written below the SNP names (e.g. T/C, T/C).

Across the locus there were 263 shared variants between the two UFM carriers with either heterozygous or homozygous differences when compared to the hg38 reference sequence. Of these 263 variants, 14 have an allele frequency <0.01 (**Figure 4a**). Both UFM carriers are heterozygous for all of the observed variants except for 2 variants in UFM2 that are homozygous for the rare allele (rs56768895 CA >C, rs828890 TT>A). None of these variants have any predicted or reported clinical significance. However, there are 3 variants which fall within the gene body of TET3, one of which (rs72818011) is in the 3’ untranslated region (3’UTR) of TET3 (**Figure 4b**). While no effect of this variant has been published, SNPs in the 3’UTR of genes have been linked with changes in mRNA stability (48,49). It is possible that this rare variant, shared by both UFM carriers, could be affecting the stability of the TET3 mRNA which could explain the lower transcription levels in these individuals. However, further analysis is needed to test whether any of the observed variants might be having an effect on the expression of TET3 or whether trans-variants or upstream genes could be driving the observed differences in TET3 expression.

### TET3 shows a different pattern of expression in mouse and human

Since humanised FXS mice fail to establish methylation of the expanded CGG repeat, we next investigated potential evolutionary changes in the expression pattern and genomic structure of the TET3 gene between human, a non-human primate and mice. A multiple sequence alignment revealed some differences in the protein sequence of all isoforms present in human, rhesus and mouse (**supplemental data**). However, none of these changes fall in regions predicted to affect the catalytic domain and the function of TET3 in any of the species (**Figure 5a**). Based on this result, it can be assumed that any differences must be in the regulation and expression of TET3 rather than in the DNA itself.

**Figure 5:**
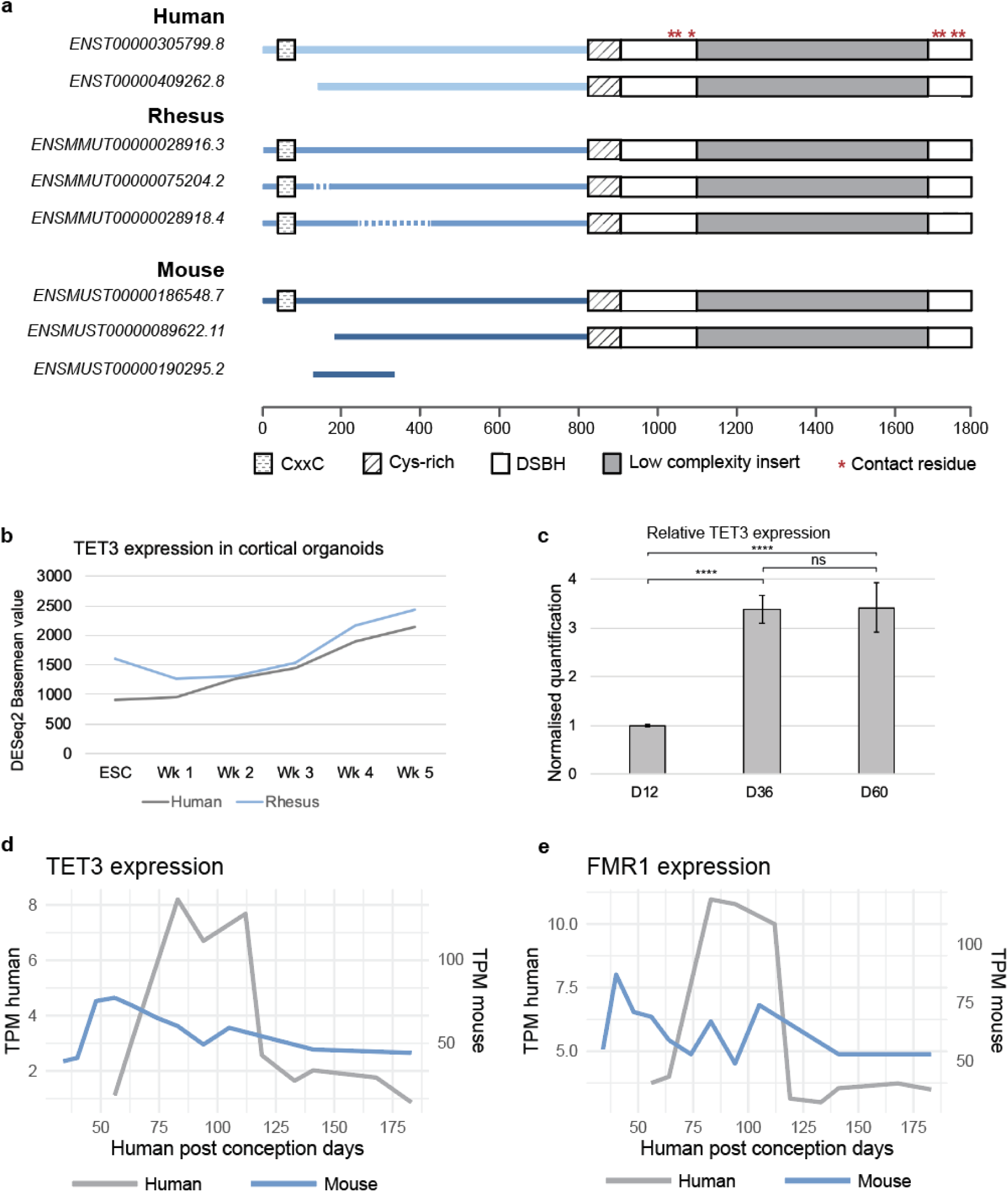
TET3 expression differences between mouse and human. a. TET3 basemean expression levels in human and rhesus cortical organoids over 5 weeks of development. b. Relative TET3 expression in cortical organoids generated from FXS iPSCs and grown over 60 days. 1-way ANOVA, Tukey’s multiple comparison test: **** p<0.001, ns not significant. c. Multiple sequence alignment of TET3 isoforms present in mouse, human and rhesus. d. TPM expression values for TET3 in the mouse and human forebrain at different age points. Note dual y-axes and scaling. e. TPM expression values for FMR1 in the human and mouse dorsolateral prefrontal cortex at different age points.

We next assessed the expression profile of TET3 in human and rhesus embryonic stem cells (ESCs) and ESC-derived neuronal cortical organoids. Expression is initially low in both human and rhesus ESCs but rises upon differentiation (**Figure 5b**). No apparent changes in the temporal expression pattern of rhesus and human TET3 was observed, suggesting that the expression dynamics of TET3 are highly conserved in primate lineages. Supportive of this data, cortical organoids derived from FXS iPSCs also show this pattern, with initial low levels of TET3 expression followed by an upregulation upon neuronal differentiation (**Figure 5c**). This upregulation can be seen to persist for at least 60 days of neuronal differentiation (**Figure 5c**) which is the point when FXS ESCs gain methylation at the FMR1 CGG repeat expansion (11). The increase in TET3 expression which coincides with the methylation time point of the CGG repeat may indicate an important role for TET3 in this process, but further validation is needed to support this hypothesis.

Finally, expression data for TET3 over time was compared between mouse and human. The translating time model was used to normalise the different gestation periods and developmental time points between the species to enable a direct inter-species comparison between developmental timepoints (45). The y-axes for the human data points were scaled to enable a comparison of the pattern of expression with the mouse samples. In the human data there is a clear peak in TET3 expression between approximately 80 and 120 post conception days (PCD) (**Figure 5d**). On the other hand, mouse TET3 expression lacks a similar peak of expression, but reaches its maximum level expression at a timepoint equivalent to 50 human PCD after which it decreases over time (**Figure 5d**). Interestingly, in human we see an increase in FMR1 expression at the same time as the peak for TET3 (**Figure 5e**). This difference in expression pattern suggests that there could be significant differences in the regulation of TET3 expression between mice and humans which may in turn affect methylation at the expanded FMR1 CGG repeat. Even though our data suggests that regulatory changes may account for the differential expression of TET3 between mice-human and UFM carriers and controls, our observations remain associative. Therefore, more research is needed to understand why UFM carriers display significantly lower levels of TET3, as a candidate gene for establishing CGG methylation in the FMR1 locus in FXS patients.

## Discussion

In this study we profiled the transcriptome of two FMR1 CGG UFM carriers. Our analysis supports and builds upon previous studies looking into transcription at the FMR1 locus and brings greater resolution to our understanding of the transcription pattern at the FMR1. We show an overall increase in expression levels in UFM carriers and high levels of a UFM cell line specific transcript deriving from the CGG repeat expansion. Our differential gene expression analysis focused on genes with a role in epigenetic modification and revealed that TET3, a gene with a role in methylation, is strongly downregulated by about 80% in two unrelated UFM carrier cell lines.

The high resolution expression profile we generated of the FMR1 locus confirms and better defines what other studies reported earlier: UFM cell lines show increased expression of FMR1-transcripts (16,18–20). We also observed a transcript originating from the CGG repeat and continuing into FMR1 intron1. This transcript is sporadically found in other transcriptomes, however, the expression of the CGG-driven transcript at high levels seems to be specific to UFM carriers. Even though only two UFM carriers were analysed, the expression level of this transcript correlates with the size of the CGG repeat expansion: in UFM2, which has a larger CGG expansion, we see higher levels than in UFM1. In FXS patient cells this transcript is not detected at all. An intron1 transcript has been observed in FXS ESCs and it has been suggested that the presence of this transcript is necessary to trigger methylation of the locus (11). This is interesting in the context of the UFM carriers as they have expression of this transcript and yet the CGG remains unmethylated. This suggests that this transcript alone is insufficient to trigger methylation at the locus. Interestingly, this phenomenon of repeat driven intronic transcription is not unique to FMR1, it is also observed in other repeat expansion disorders where repeat driven transcription is often a pathogenic mechanism (50,51).

Based on our analyses, TET3 is a strong candidate for involvement in FMR1 methylation. It is strongly downregulated in the UFM carriers and becomes highly expressed in the window when the expanded FMR1 repeat gains methylation. Although TET3 is also highly conserved in mice, the expression patterns across developmental time differ between the two species which hints towards species-specific regulatory properties which could underlie subtle differences in function. A previous exome comparative analysis on UFM, FXS and control neither find any significant mutations in the epigenetic machinery genes nor in the proximal regulatory sequences of FMR1 (18). Our analysis of WGS data from both UFM individuals reveal a variant in the 3’UTR of TET3 that could have an effect on mRNA stability levels, but further testing is required to see if this is the case. In theory, it is possible that a healthy individual without the FMR1 CGG repeat expansion could also carry the variant that we seek in the UFM carriers. In this instance it would be difficult to find the causal mutation when comparing to the reference sequence.

The TET enzymes are a family of methylcytosine dioxygenases. TET3 consists of a CXXC domain which recognises and binds to specific GC-rich DNA sequences and also a catalytic domain which plays a key role in the establishment of the 5hydroxymethylated cytosine (5hmC) DNA modification by converting 5mC to 5hmC (52). One possible mechanism by which TET3 influences methylation at the FMR1 CGG repeat expansion Is through the establishment of 5hmC, which was initially thought to be a transitional state of DNA demethylation. Recent studies show, however, that 5hmC is also a stable epigenetic mark which is enriched in the brain (52,53). Interestingly, 5hmC has also been shown to be enriched around the CGG repeat in the neurons of FXS patients (54). Therefore, It is possible that TET3 plays a crucial role in the establishment of 5hmC (and possibly 5mC) at the expanded FMR1 locus. Even though the exact mechanism remains elusive, lower levels of TET3 could lead to a reduction or lack of DNA methylation at the FMR1 promoter in UFM carriers (**Figure 6**). Out of all three TET enzymes, TET1, TET2 and TET3, the role of TET3 remains obscure. While TET1 and TET2 have been reported to be important for DNA demethylation, there are several observations that suggest that TET3 may have a different role. Cell lines with a TET3 knockout have been shown to have a global decrease in methylation levels, specifically at gene promoters (55,56). This is in stark contrast to an assumed role of TET enzymes in DNA demethylation, where the opposite would be expected. Another study comparing TET3 knockout mouse embryonic stem cells (mESCs) with mESCs that contained a version of TET3 with a non-functioning catalytic domain found that in both conditions, differentially methylated regions were hypomethylated suggesting that TET3 is required for methylation at specific gene promoters. This study found that DNA methyltransferase 1 (DNMT1) is a direct target of TET3 and that without TET3, DNMT1 was downregulated (56). This observation offers an alternate, or perhaps complementary mechanism through which TET3 could operate. Reduced levels of TET3 result in a downregulation and lack of sufficient DNMT1 which is needed to methylate the CGG expansion (**Figure 6**). Taken together, the studies in TET3 Knock out mice suggest that as opposed to the other TET enzymes, TET3 may be required for the establishment of DNA methylation, a distinct functional characteristic that is in agreement with the observed deficits of TET3 expression in UFM carriers.

**Fig 6:**
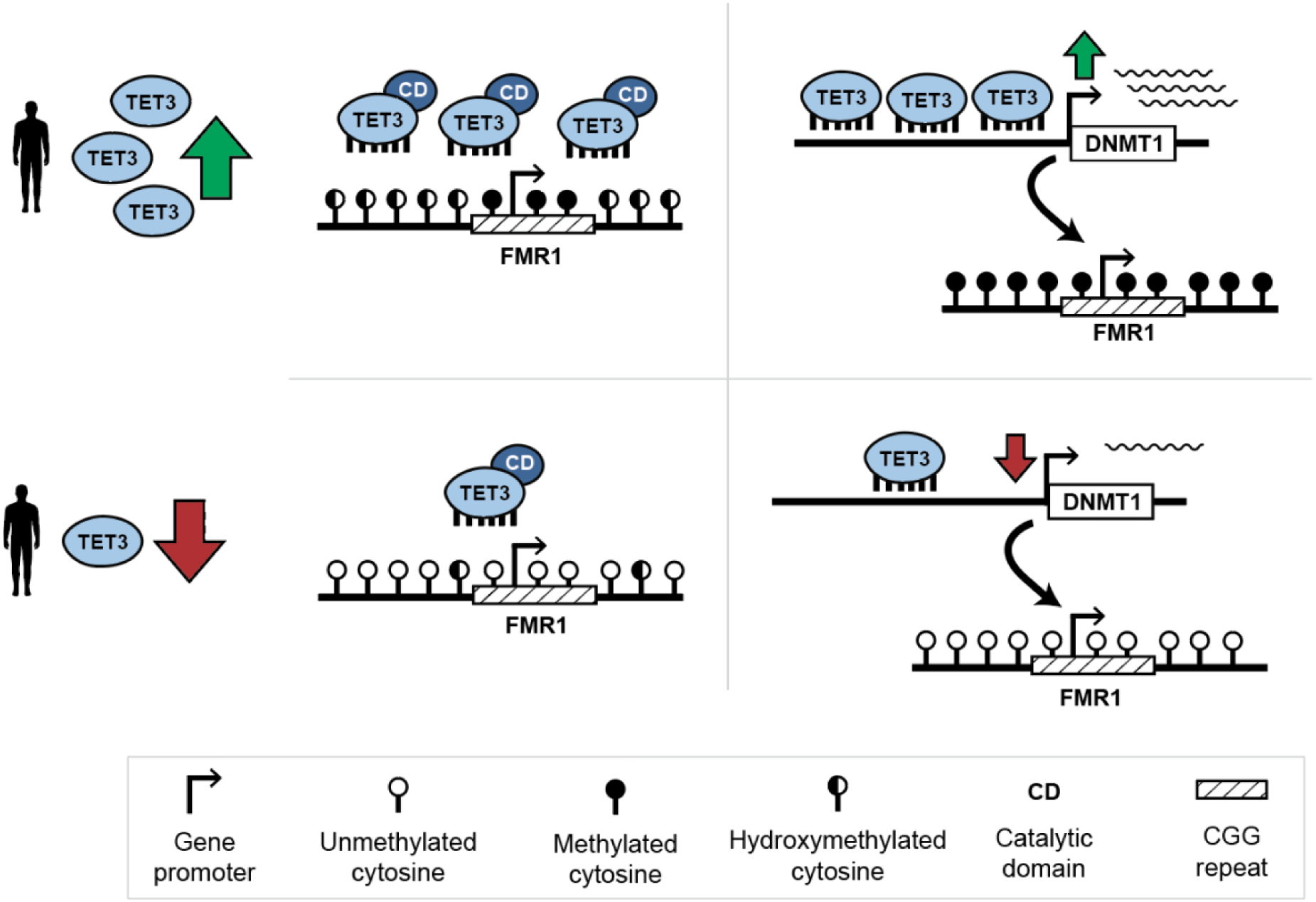
Potential model for the role of TET3 in FMR1 repeat methylation. a. In individuals with higher levels of TET3, the enzyme potentially functions in both a catalytic and non-catalytic role to affect methylation levels at the FMR1 CGG repeat expansion. Utilising the catalytic domain, TET3 may be directly responsible for the establishment of 5hmC at the locus. In a non-catalytic function, TET3 directly regulates DNMT1 expression which in turn is responsible for methylation at the FMR1 CGG repeat expansion. b. In individuals with reduced TET3 levels, there is insufficient enzyme to establish and maintain 5hmC at the FMR1 promoter. The lower levels of TET3 also affect DNMT1 expression leading to a depletion in DNMT1 activity and a lack of methylation at the FMR1 locus.

The nature and timing of methylation at the FMR1 locus raises many difficulties in the study of methylation dynamics at the expanded CGG repeat. There are very few cell lines where this is possible. Naïve reversion of FXS iPSCs has been shown to reactivate expression at the locus and remove methylation (57). However, this has been shown to be a result of contraction of the CGG repeat and remethylation does not occur upon differentiation back to primed cells. In addition, the instability of the CGG repeat makes iPSCs derived from UFM carriers unreliable model systems as in some cases they have been shown to gain methylation after iPSC conversion (19,20). Despite these limitations, we believe that our identification of TET3 as a candidate in the role of FMR1 CGG repeat methylation is robust and a novel finding in the field which has the potential to deepen our understanding of the methylation dynamics of the CGG repeat in the FMR1 locus.

## Conclusions

Our study offers a unique insight into the transcriptomes and genomes of two UFM carriers. We show a high resolution overview of transcription at FMR1 locus, highlighting intronic transcripts expressed in high levels specifically in the UFM carriers. Our differential gene expression analysis shows a strong downregulation of ~80% for TET3 in two unrelated UFM carriers when compared to controls. Both UFM carriers have a variant in the TET3 3’UTR that has the potential to influence mRNA stability and could account for the observed differential expression. Loss or knockdown of TET3 has been associated with loss of DNA methylation particularly at gene promoters, similar to the location of the CGG repeat in the FMR1 locus. These effects have been linked to both the catalytic function of TET3 in converting 5mC to 5hmC and a non-catalytic role of the enzyme binding to specific promoters. Taken together, the findings described in this study put forward TET3 as a prime candidate for mediating the methylation of the CGG repeat expansion in FXS. While further research is needed, the identification of an epigenetic factor responsible for the silencing of the FMR1 locus may have important consequences for our understanding of FXS as well as the development of future therapeutic strategies.

## Supporting information

Full MSA of TET3

## List of abbreviations

3’UTR: Three prime untranslated region
5’UTR: Five prime untranslated region
DNMT1: DNA methyltransferase 1
ESC: Embryonic stem cell
EZH2: Enhancer Of Zeste 2 Polycomb Repressive Complex 2 Subunit
FMR1: Fragile X Messenger Ribonucleoprotein 1
FMRP: Fragile X Messenger Ribonucleoprotein
FXS: Fragile X syndrome
FXTAS: Fragile X-associated tremor/ataxia syndrome
iPSC: Induced pluripotent stem cell
mESCs: Mouse embryonic stem cells
PCD: Post conception days
PM: Pre-mutation
FXPOI: Fragile X-Associated Primary Ovarian Insufficiency
SETD1B: SET Domain Containing 1B
TET3: Tet methylcytosine dioxygenase 3
UFM: Unmethylated full mutation
WGS: Whole genome sequencing

## Declarations

### Ethics approval and consent to participate

All procedures performed in studies involving human participants were in accordance with the ethical standards of the institutional and/or national research committee and with the 1964 Helsinki declaration and its later amendments or comparable ethical standards.

### Consent for publication

All authors consent to the publication of this manuscript. Written informed consent was obtained from the participants (or their legal guardians) for publication of any potentially identifiable data included in this article.

### Availability of data and materials

RNA and whole genome sequencing data have been deposited at the European Genome-Phenome Archive (EGA, https://ega-archive.org), under accession numbers EGAS50000000647 (RNA-seq) and EGAS50000000648 (WGS). The data are accessible to the FXS research community via the controlled access procedure of the EGA.

### Authors’ contributions

Conceptualization, G.F., F.M.J.J., E.T. and R.W.; Methodology, G.F., F.M.J.J.; Investigation & Validation, G.F., V.B.; Data Curation, G.F., F.M.J.J.; Writing – Original Draft, G.F and F.M.J.J.; Writing – Review & Editing; F.M.J.J., G.F., E.T., R.W.; Visualization, G.F.; Supervision, F.M.J.J.; Project Administration, F.M.J.J.; Funding Acquisition, F.M.J.J.

## Acknowledgements

We would like to acknowledge the SILS MAD sequencing group for technical assistance with Illumina sequencing; We like to thank Karen Usdin, the Evolutionary Neurogenomics Group, the Molecular Neuroscience group, and others at the Swammerdam Institute for Life Sciences for helpful discussions. This project was funded by an ERC starting grant to F.M.J.J

