## Supplemental figures for "Transcriptomic profiling of unmethylated full mutation carriers implicates TET3 in FMR1 CGG repeat expansion methylation dynamics in Fragile X syndrome"

| Name | Function |
|---|---|
| DNMT1 | DNA-methyltransferase |
| DNMT3a | DNA-methyltransferase |
| DNMT3b | DNA-methyltransferase |
| EZH1 | Methyltransferase (histone) |
| EZH2 | Methyltransferase (histone) |
| G9a-GLP | Methyltransferase (histone) |
| JARID1A/KDM5A | demethylase (histone) |
| JARID1B/KDM5B | demethylase (histone) |
| JARID1C/KDM5C | demethylase (histone) |
| JARID1D/KDM5D | demethylase (histone) |
| JHDM2A/KDM3A | demethylase (histone) |
| JHDM2B//KDM3B | demethylase (histone) |
| JHDM2C/KDM3C | demethylase (histone) |
| JHDM3A/KDM4A | demethylase (histone) |
| JHDM3B/KDM4B | demethylase (histone) |
| JHDM3C/KDM4C | demethylase (histone) |
| JHDM3D/KDM4D | demethylase (histone) |
| JMJD3/KDM6B | demethylase (histone) |
| KIAA1718/KDM7A | demethylase (histone) |
| KMT1D/EHMT1 | Methyltransferase (histone) |
| LSD1/KDM1A | demethylase (histone) |
| LSD2/KDM1B | demethylase (histone) |
| MLL1/KMT2A | Methyltransferase (histone) |
| MLL2/KMT2B | Methyltransferase (histone) |
| MLL3/KMT2C | Methyltransferase (histone) |
| MLL4/KMT2D | Methyltransferase (histone) |
| NO66/MAPJD | demethylase (histone) |
| PHF2 | demethylase (histone) |
| PHF8 | demethylase (histone) |
| PHF8 | demethylase (histone) |
| PRDM9 | Methyltransferase (histone) |
| SET1A | Methyltransferase (histone) |
| SET1B | Methyltransferase (histone) |
| SET7 | Methyltransferase (histone) |
| SETDB1 | Methyltransferase (histone) |
| SMYD1/KMT3D | Methyltransferase (histone) |
| SMYD2/KMT3C | Methyltransferase (histone) |
| SUV39H1 | Methyltransferase (histone) |
| SUV39H2 | Methyltransferase (histone) |
| TET1 | Methylcytosine dioxygenase |
| TET2 | Methylcytosine dioxygenase |
| TET3 | Methylcytosine dioxygenase |
| UTX/KDM6A | demethylase (histone) |
| UTY/KDM6C | demethylase (histone) |

***Supplemental table 1***

*List of 44 candidate genes with roles in epigenetic modification of DNA*

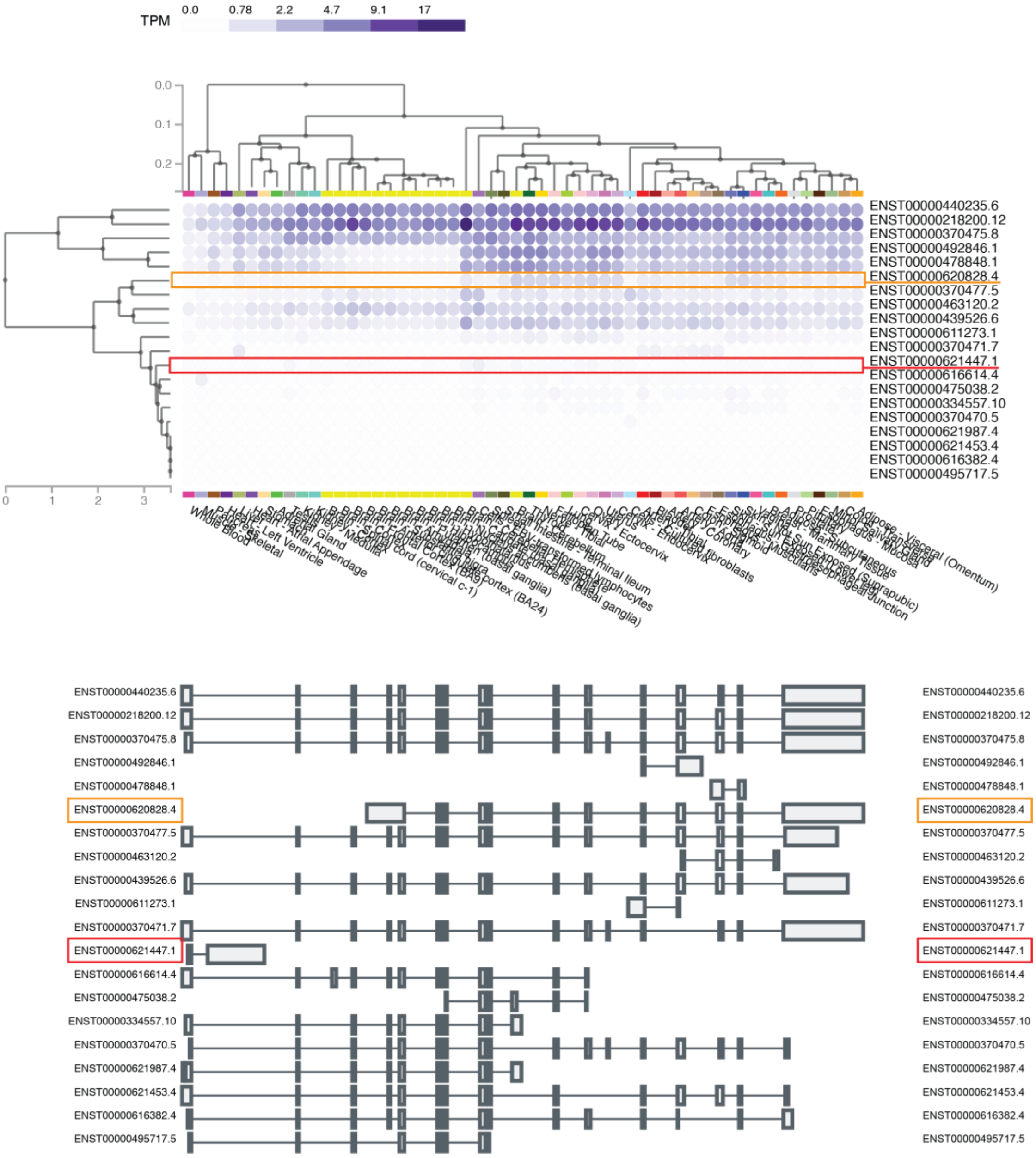


***Supplemental figure 1***

*FMR1 transcript information from gTEX*

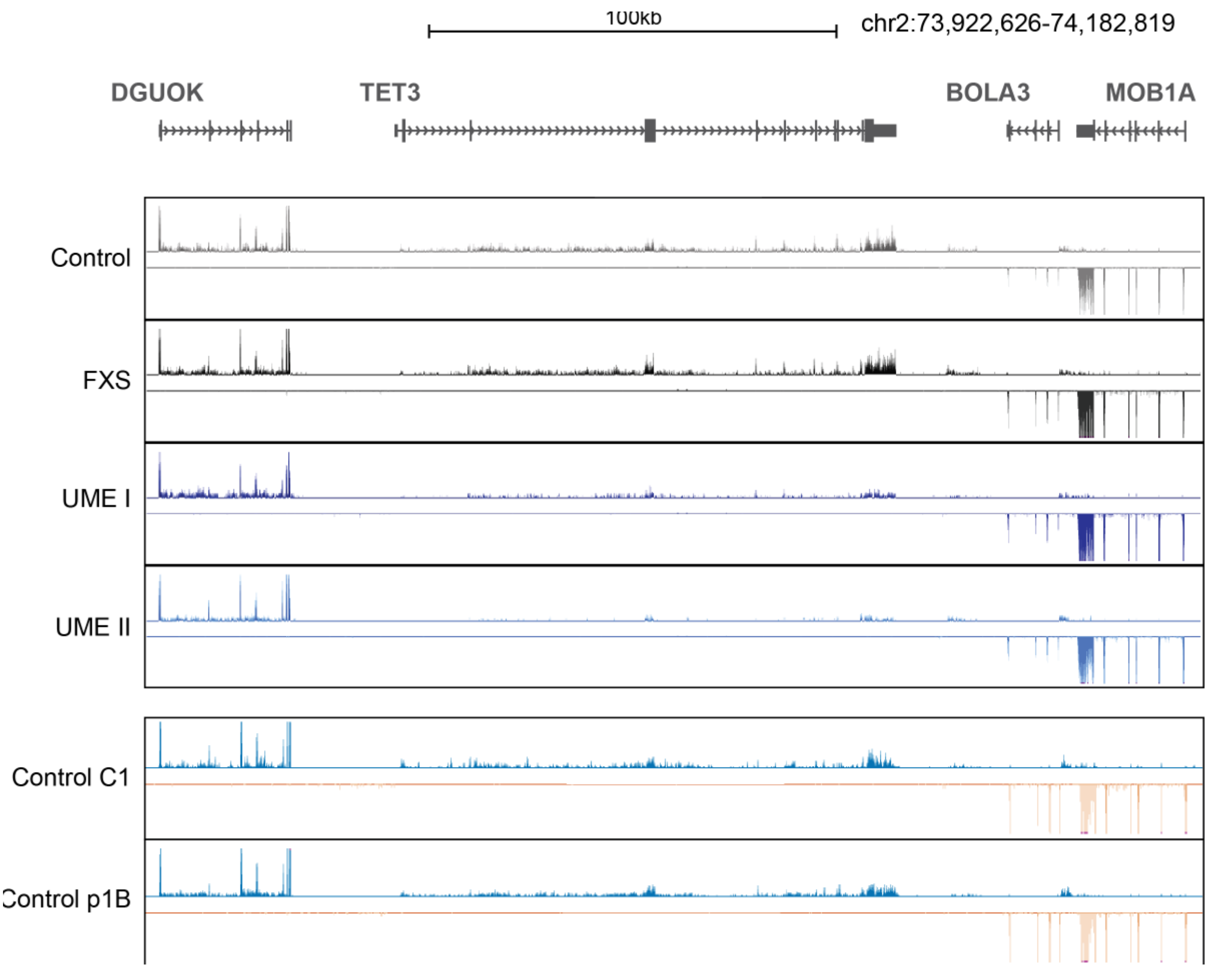


***Supplemental figure 2***

*Coverage plots of RNA-seq data at the TET3 gene from fibroblasts obtained from a control individual, an FXS patient and two FMR1 UFM carriers. Coverage tracks are scaled based on the total number of reads successfully mapped to the genome. Coverage tracks were generated from track collections and show merged tracks from 3 replicates. Control C1 and p1B showing RNA-seq coverage from unrelated fibroblast cell lines generated by Fernandes et al (jn prep) scaled to neighbouring genes.*
