## Supplementary material for "Transcriptomic profiling of unmethylated full mutation carriers implicates TET3 in FMR1 CGG repeat expansion methylation dynamics in Fragile X syndrome": Full MSA of TET3

CxxC domain

```
CLUSTAL O(1.2.4) multiple sequence alignment


Mouse_ENSMUST00000186548.7        MSQFQVPLAVQPDLSGLYDFPQGQVMVGGFQGPGLPMAGSETQLRGGGDGRKKRKRCGTC 60
Human_ENST00000409262.8           MSQFQVPLAVQPDLPGLYDFPQRQVMVGSFPGSGLSMAGSESQLRGGGDGRKKRKRCGTC 60
Rhesus_ENSMMUT00000028918.4       MSQFQVPLAVQPDLPGLYDFPQGQVMVGGFPGPELSMAGSESQLRGGGDGRKKRKRCGTC 60
Rhesus_ENSMMUT00000028916.3       MSQFQVPLAVQPDLPGLYDFPQGQVMVGGFPGPELSMAGSESQLRGGGDGRKKRKRCGTC 60
Rhesus_ENSMMUT00000075204.2       MSQFQVPLAVQPDLPGLYDFPQGQVMVGGFPGPELSMAGSESQLRGGGDGRKKRKRCGTC 60
                                  ************** ******* *****.* *  * *****:******************

Mouse_ENSMUST00000186548.7        DPCRRLENCGSCTSCTNRRTHQICKLRKCEVLKKKAGLLKEVEINAREGTGPWAQGATVK 120
Human_ENST00000409262.8           EPCRRLENCGACTSCTNRRTHQICKLRKCEVLKKKVGLLKEVEIKAGEGAGPWGQGAAVK 120
Rhesus_ENSMMUT00000028918.4       EPCRRLENCGACTSCTNRRTHQICKLRKCEVLKKKVGLLKEVEIKAGEGAGPWGQGAAVK 120
Rhesus_ENSMMUT00000028916.3       EPCRRLENCGACTSCTNRRTHQICKLRKCEVLKKKVGLLKEVEIKAGEGAGPWGQGAAVK 120
Rhesus_ENSMMUT00000075204.2       EPCRRLENCGACTSCTNRRTHQICKLRKCEVLKKKVGLLKE------------------- 101
                                  :*********:************************.*****

Mouse_ENSMUST00000186548.7        TGSELSPVDGPVPGQMDSGPVYHGDSRQLSTSGAPVNGAREPAGPGLLGAAGPWRVDQKP 180
Human_ENST00000409262.8           TGSELSPVDGPVPGQMDSGPVYHGDSRQLSASGVPVNGAREPAGPSLLGTGGPWRVDQKP 180
Rhesus_ENSMMUT00000028918.4       TGSELSPVDGPVPGQMDSGPVYHGDSRQLSASGVPVNGAREPAGPSLLGTGGPWRVDQKP 180
Rhesus_ENSMMUT00000028916.3       TGSELSPVDGPVPGQMDSGPVYHGDSRQLSASGVPVNGAREPAGPSLLGTGGPWRVDQKP 180
Rhesus_ENSMMUT00000075204.2       TGSELSPVDGPVPGQMDSGPVYHGDSRQLSASGVPVNGAREPAGPSLLGTGGPWRVDQKP 161
                                  ******************************:**.***********.***:.*********

Mouse_ENSMUST00000186548.7        DWEAASGPTHAARLEDAHDLVAFSAVAEAVSSYGALSTRLYETFNREMSREAGSNGRGPR 240
Human_ENST00000409262.8           DWEAAPGPAHTARLEDAHDLVAFSAVAEAVSSYGALSTRLYETFNREMSREAGNNSRGPR 240
Rhesus_ENSMMUT00000028918.4       DWEAAPGPAHTARLEDAHDLVAFSAVAEAVSSYGALSTRLYETFNREMSREAGNNSRGSR 240
Rhesus_ENSMMUT00000028916.3       DWEAAPGPAHTARLEDAHDLVAFSAVAEAVSSYGALSTRLYETFNREMSREAGNNSRGSR 240
Rhesus_ENSMMUT00000075204.2       DWEAAPGPAHTARLEDAHDLVAFSAVAEAVSSYGALSTRLYETFNREMSREAGNNSRGSR 221
                                  ***** **:*:******************************************.*.** *

Mouse_ENSMUST00000186548.7        --PESCSEGSEDLDTLQTALALARHGMKPPNCTCDGPECPDFLEWLEGKIKSMAMEGGQG 298
Human_ENST00000409262.8           PGPEGCSAGSEDLDTLQTALALARHGMKPPNCNCDGPECPDYLEWLEGKIKSVVMEGGEE 300
Rhesus_ENSMMUT00000028918.4       PGPEGCSAGSEDLDTLQTALALARH----------------------------------- 265
Rhesus_ENSMMUT00000028916.3       PGPEGCSAGSEDLDTLQTALALARHGMKPPNCNCDGPECPDYLEWLEGKIKSVVMEGGEE 300
Rhesus_ENSMMUT00000075204.2       PGPEGCSAGSEDLDTLQTALALARHGMKPPNCNCDGPECPDYLEWLEGKIKSVVMEGGEE 281
                                    **.** *****************

Mouse_ENSMUST00000186548.7        RPRLPGALPPSEAGLPAPSTRPPLLSSEVPQVPPLEGLPLSQSALSIAKEKNISLQTAIA 358
Human_ENST00000409262.8           RPRLPGPLPPGEAGLPAPSTRP-LLSSEVPQISPQEGLPLSQSALSIAKEKNISLQTAIA 359
Rhesus_ENSMMUT00000028918.4       ------------------------------------------------------------ 265
Rhesus_ENSMMUT00000028916.3       RPRLPGPLPPGEAGLPAPSTRP-LLSSEVPQISPQEGLPLSQSALSIAKEKNISLQTAIA 359
Rhesus_ENSMMUT00000075204.2       RPRLPGPLPPGEAGLPAPSTRP-LLSSEVPQISPQEGLPLSQSALSIAKEKNISLQTAIA 340


Mouse_ENSMUST00000186548.7        IEALTQLSSALPQPSHSTSQASCPLPEALSPSAPFRSPQSYLRAPSWPVVPPEEHPSFAP 418
Human_ENST00000409262.8           IEALTQLSSALPQPSHSTPQASCPLPEALSPPAPFRSPQSYLRAPSWPVVPPEEHSSFAP 419
Rhesus_ENSMMUT00000028918.4       ------------------------------------------------------------ 265
Rhesus_ENSMMUT00000028916.3       IEALTQLSSALPQPSHSTPQASCPLPEALSPPAPFRSPQSYLRAPSWPVVPPEEHSSFAP 419
Rhesus_ENSMMUT00000075204.2       IEALTQLSSALPQPSHSTPQASCPLPEALSPPAPFRSPQSYLRAPSWPVVPPEEHSSFAP 400


Mouse_ENSMUST00000186548.7        DSPAFPPATPRPEFSEAWGTDTPPATPRNSWPVPRPSPDPMAELEQLLGSASDYIQSVFK 478
Human_ENST00000409262.8           DSSAFPPATPRTEFPEAWGTDTPPATPRSSWPMPRPSPDPMAELEQLLGSASDYIQSVFK 479
Rhesus_ENSMMUT00000028918.4       -------ATPRTEFPEVWGTDTPPATPRSSWPMPRPSHDPMAELEQLLGSASDYIQSVFK 318
Rhesus_ENSMMUT00000028916.3       DSSAFPPATPRTEFPEVWGTDTPPATPRSSWPMPRPSHDPMAELEQLLGSASDYIQSVFK 479
Rhesus_ENSMMUT00000075204.2       DSSAFPPATPRTEFPEVWGTDTPPATPRSSWPMPRPSHDPMAELEQLLGSASDYIQSVFK 460
                                         **** ** *.***********.***:**** **********************

Mouse_ENSMUST00000186548.7        RPEALPTKPKVKVEAPSSSPAPVPSPISQREAPLLSSEPDTHQKAQTALQQHLHHKRNLF 538
Human_ENST00000409262.8           RPEALPTKPKVKVEAPSSSPAPAPSPVLQREAPTPSSEPDTHQKAQTALQQHLHHKRSLF 539
Rhesus_ENSMMUT00000028918.4       RPEALPTKPKVKVEAPSSSPALAPSPVLQREAPTPSSEPDTHQKAQTALQQHLHHKRSLF 378
Rhesus_ENSMMUT00000028916.3       RPEALPTKPKVKVEAPSSSPALAPSPVLQREAPTPSSEPDTHQKAQTALQQHLHHKRSLF 539
Rhesus_ENSMMUT00000075204.2       RPEALPTKPKVKVEAPSSSPALAPSPVLQREAPTPSSEPDTHQKAQTALQQHLHHKRSLF 520
                                  ********************* .***: *****  **********************.**

Mouse_ENSMUST00000186548.7        LEQAQDASFPTSTEPQAPGWWAPPGSPAPRPPDKPPKEKKKKPPTPAGGPVGAEKTTPGI 598
Human_ENST00000409262.8           LEQVHDTSFPAPSEPSAPGWWPPPSSPVPRLPDRPPKEKKKKLPTPAGGPVGTEKAAPGI 599
Rhesus_ENSMMUT00000028918.4       LEQAHDTSFPAPSEPSAPGWWPPPSSPAPRLPDRPPKEKKKKLPTPAGGPVGTEKAAPGI 438
Rhesus_ENSMMUT00000028916.3       LEQAHDTSFPAPSEPSAPGWWPPPSSPAPRLPDRPPKEKKKKLPTPAGGPVGTEKAAPGI 599
Rhesus_ENSMMUT00000075204.2       LEQAHDTSFPAPSEPSAPGWWPPPSSPAPRLPDRPPKEKKKKLPTPAGGPVGTEKAAPGI 580
                                  ***.:*:***: :**.***** **.**.** **:******** *********:**::***
```

```
Mouse_ENSMUST00000186548.7       KTSVRKPIQIKKSRSRDMQPLFLPVRQIVLEGLKPQASEGQAPLPAQLSVPPPASQGAAS     658
Human_ENST00000409262.8          KPSVRKPIQIKKSRPREAQPLFFPVRQIVLEGLRSPASQEVQAHP---PAPL-----PAS     651
Rhesus_ENSMMUT00000028918.4      KPSVRKPIQIKKSRPREAQPLFFPVRQIVLEGLRSPASQEVQAHP---PAPL-----PAS     490
Rhesus_ENSMMUT00000028916.3      KPSVRKPIQIKKSRPREAQPLFFPVRQIVLEGLRSPASQEVQAHP---PAPL-----PAS     651
Rhesus_ENSMMUT00000075204.2      KPSVRKPIQIKKSRPREAQPLFFPVRQIVLEGLRSPASQEVQAHP---PAPL-----PAS     632
                                 * ************ *: **** **********:  **:     *    .*       **

Mouse_ENSMUST00000186548.7       QSCATPLTPEPSLALFAPSPSGDSLLPPTQEMRSPSPMVALQSGSTGGPLPPADDKLEEL     718
Human_ENST00000409262.8          QGSAVPLPPEPSLALFAPSPSRDSLLPPTQEMRSPSPMTALQPGST-GPLPPADDKLEEL     710
Rhesus_ENSMMUT00000028918.4      QGSAVPLPPEPSLALFAPSPSRDSLLPPTQEMRSPSPMTTLQPGST-GPLPPADDKLEEL     549
Rhesus_ENSMMUT00000028916.3      QGSAVPLPPEPSLALFAPSPSRDSLLPPTQEMRSPSPMTTLQPGST-GPLPPADDKLEEL     710
Rhesus_ENSMMUT00000075204.2      QGSAVPLPPEPSLALFAPSPSRDSLLPPTQEMRSPSPMTTLQPGST-GPLPPADDKLEEL     691
                                 *..*.** ************* ****************.:** *** *************

Mouse_ENSMUST00000186548.7       IRQFEAEFGDSFGLPGPPSVPIQEPENQSTCLPAPESPFATRSPKKIKIESSGAVTVLST     778
Human_ENST00000409262.8          IRQFEAEFGDSFGLPGPPSVPIQDPENQQTCLPAPESPFATRSPKQIKIESSGAVTVLST     770
Rhesus_ENSMMUT00000028918.4      IRQFEAEFGDSFGLPGPPSVPIQDPENQQTCLPAPESPFATRSPKQIKIESSGAVTVLST     609
Rhesus_ENSMMUT00000028916.3      IRQFEAEFGDSFGLPGPPSVPIQDPENQQTCLPAPESPFATRSPKQIKIESSGAVTVLST     770
Rhesus_ENSMMUT00000075204.2      IRQFEAEFGDSFGLPGPPSVPIQDPENQQTCLPAPESPFATRSPKQIKIESSGAVTVLST     751
                                 ***********************:****.****************:**************

Mouse_ENSMUST00000186548.7       TCFHSEEGGQEATPTKAENPLTPTLSGFLESPLKYLDTPTKSLLDTPAKKAQSEFPTCDC     838
Human_ENST00000409262.8          TCFHSEEGGQEATPTKAENPLTPTLSGFLESPLKYLDTPTKSLLDTPAKRAQAEFPTCDC     830
Rhesus_ENSMMUT00000028918.4      TCFHSEEGGQEATPTKAENPLTPTLSGFLESPLKYLDTPTKSLLDTPAKRAQAEFPTCDC     669
Rhesus_ENSMMUT00000028916.3      TCFHSEEGGQEATPTKAENPLTPTLSGFLESPLKYLDTPTKSLLDTPAKRAQAEFPTCDC     830
Rhesus_ENSMMUT00000075204.2      TCFHSEEGGQEATPTKAENPLTPTLSGFLESPLKYLDTPTKSLLDTPAKRAQAEFPTCDC     811
                                 *************************************************:**:*******

Mouse_ENSMUST00000186548.7       VEQIVEKDEGPYYTHLGSGPTVASIRELMEDRYGEKGKAIRIEKVIYTGKEGKSSRGCPI     898
Human_ENST00000409262.8          VEQIVEKDEGPYYTHLGSGPTVASIRELMEERYGEKGKAIRIEKVIYTGKEGKSSRGCPI     890
Rhesus_ENSMMUT00000028918.4      VEQIVEKDEGPYYTHLGSGPTVASIRELMEERYGEKGKAIRIEKVIYTGKEGKSSRGCPI     729
Rhesus_ENSMMUT00000028916.3      VEQIVEKDEGPYYTHLGSGPTVASIRELMEERYGEKGKAIRIEKVIYTGKEGKSSRGCPI     890
Rhesus_ENSMMUT00000075204.2      VEQIVEKDEGPYYTHLGSGPTVASIRELMEERYGEKGKAIRIEKVIYTGKEGKSSRGCPI     871
                                 ******************************:*****************************

Mouse_ENSMUST00000186548.7       AKWVIRRHTLEEKLLCLVRHRAGHHCQNAVIVILILAWEGIPRSLGDTLYQELTDTLRKY     958
Human_ENST00000409262.8          AKWVIRRHTLEEKLLCLVRHRAGHHCQNAVIVILILAWEGIPRSLGDTLYQELTDTLRKY     950
Rhesus_ENSMMUT00000028918.4      AKWVIRRHTLEEKLLCLVRHRAGHHCQNAVIVILILAWEGIPRSLGDTLYQELTDTLRKY     789
Rhesus_ENSMMUT00000028916.3      AKWVIRRHTLEEKLLCLVRHRAGHHCQNAVIVILILAWEGIPRSLGDTLYQELTDTLRKY     950
Rhesus_ENSMMUT00000075204.2      AKWVIRRHTLEEKLLCLVRHRAGHHCQNAVIVILILAWEGIPRSLGDTLYQELTDTLRKY     931
                                 ************************************************************

Mouse_ENSMUST00000186548.7       GNPTSRRCGLNDDRTCACQGKDPNTCGASFSFGCSWSMYFNGCKYARSKTPRKFRLTGDN     1018
Human_ENST00000409262.8          GNPTSRRCGLNDDRTCACQGKDPNTCGASFSFGCSWSMYFNGCKYARSKTPRKFRLAGDN     1010
Rhesus_ENSMMUT00000028918.4      GNPTSRRCGLNDDRTCACQGKDPNTCGASFSFGCSWSMYFNGCKYARSKTPRKFRLAGDN     849
Rhesus_ENSMMUT00000028916.3      GNPTSRRCGLNDDRTCACQGKDPNTCGASFSFGCSWSMYFNGCKYARSKTPRKFRLAGDN     1010
Rhesus_ENSMMUT00000075204.2      GNPTSRRCGLNDDRTCACQGKDPNTCGASFSFGCSWSMYFNGCKYARSKTPRKFRLAGDN     991
                                 ********************************************************:***

Mouse_ENSMUST00000186548.7       PKEEEVLRNSFQDLATEVAPLYKRLAPQAYQNQVTNEDVAIDCRLGLKEGRPFSGVTACM     1078
Human_ENST00000409262.8          PKEEEVLRKSFQDLATEVAPLYKRLAPQAYQNQVTNEEIAIDCRLGLKEGRPFAGVTACM     1070
Rhesus_ENSMMUT00000028918.4      PKEEEVLRKSFQDLATEVAPLYKRLAPQAYQNQVTNEEIAIDCRLGLKEGRPFAGVTACM     909
Rhesus_ENSMMUT00000028916.3      PKEEEVLRKSFQDLATEVAPLYKRLAPQAYQNQVTNEEIAIDCRLGLKEGRPFAGVTACM     1070
Rhesus_ENSMMUT00000075204.2      PKEEEVLRKSFQDLATEVAPLYKRLAPQAYQNQVTNEEIAIDCRLGLKEGRPFAGVTACM     1051
                                 ********:****************************::**************:******

Mouse_ENSMUST00000186548.7       DFCAHAHKDQHNLYNGCTVVCTLTKEDNRCVGQIPEDEQLHVLPLYKMASTDEFGSEENQ     1138
Human_ENST00000409262.8          DFCAHAHKDQHNLYNGCTVVCTLTKEDNRCVGKIPEDEQLHVLPLYKMANTDEFGSEENQ     1130
Rhesus_ENSMMUT00000028918.4      DFCAHAHKDQHNLYNGCTVVCTLTKEDNRCVGKIPEDEQLHVLPLYKMASTDEFGSEENQ     969
Rhesus_ENSMMUT00000028916.3      DFCAHAHKDQHNLYNGCTVVCTLTKEDNRCVGKIPEDEQLHVLPLYKMASTDEFGSEENQ     1130
Rhesus_ENSMMUT00000075204.2      DFCAHAHKDQHNLYNGCTVVCTLTKEDNRCVGKIPEDEQLHVLPLYKMASTDEFGSEENQ     1111
                                 ********************************:****************.**********

Mouse_ENSMUST00000186548.7       NAKVSSGAIQVLTAFPREVRRLPEPAKSCRQRQLEARKAAAEKKKLQKEKLSTPEKIKQE     1198
Human_ENST00000409262.8          NAKVGSGAIQVLTAFPREVRRLPEPAKSCRQRQLEARKAAAEKKKIQKEKLSTPEKIKQE     1190
Rhesus_ENSMMUT00000028918.4      NAKVGSGAIQVLTAFPREVRRLPEPAKSCRQRQLEARKAAAEKKKIQKEKLSTPEKIKQE     1029
Rhesus_ENSMMUT00000028916.3      NAKVGSGAIQVLTAFPREVRRLPEPAKSCRQRQLEARKAAAEKKKIQKEKLSTPEKIKQE     1190
Rhesus_ENSMMUT00000075204.2      NAKVGSGAIQVLTAFPREVRRLPEPAKSCRQRQLEARKAAAEKKKIQKEKLSTPEKIKQE     1171
                                 ****.***************************************:***************

Mouse_ENSMUST00000186548.7       ALELAGVTTDPGLSLKGGLSQQSLKPSLKVEPQNHFSSFKYSGNAVVESYSVLGSCRPSD     1258
Human_ENST00000409262.8          ALELAGITSDPGLSLKGGLSQQGLKPSLKVEPQNHFSSFKYSGNAVVESYSVLGNCRPSD     1250
Rhesus_ENSMMUT00000028918.4      ALELAGVTSDPGLSLKGGLSQQGLKPSLKVEPQNHFSSFKYSSNAVVESYSVLGNCRPSD     1089
Rhesus_ENSMMUT00000028916.3      ALELAGVTSDPGLSLKGGLSQQGLKPSLKVEPQNHFSSFKYSSNAVVESYSVLGNCRPSD     1250
Rhesus_ENSMMUT00000075204.2      ALELAGVTSDPGLSLKGGLSQQGLKPSLKVEPQNHFSSFKYSSNAVVESYSVLGNCRPSD     1231
                                 ******:*:*************.*******************.***********.*****

Mouse_ENSMUST00000186548.7       PYSMSSVYSYHSRYAQPGLASVNGFHSKYTLPSFGYYGFPSSNPVFPSQFLGPSAWGHGG     1318
Human_ENST00000409262.8          PYSMNSVYSYHSYYAQPSLTSVNGFHSKYALPSFSYYGFPSSNPVFPSQFLGPGAWGHSG     1310
Rhesus_ENSMMUT00000028918.4      PYSMNSVYSYHSYYAQPSLTSVNGFHSKYALPSFGYYGFPSSNPVFPSQFLGPGAWGHSG     1149
Rhesus_ENSMMUT00000028916.3      PYSMNSVYSYHSYYAQPSLTSVNGFHSKYALPSFGYYGFPSSNPVFPSQFLGPGAWGHSG     1310
Rhesus_ENSMMUT00000075204.2      PYSMNSVYSYHSYYAQPSLTSVNGFHSKYALPSFGYYGFPSSNPVFPSQFLGPGAWGHSG     1291
                                 ****.******* ****.*:*********:****.******************.****.*
```

CYS rich | Catalytic residue | Large low complexity insert

```
Mouse_ENSMUST00000186548.7      SGGSFEKKPDLHALHNSLNPAYGGAEFAELPGQAVATDNHHPIPHHQQPAYPGPKEYLLP     1378
Human_ENST00000409262.8         SSGSFEKKPDLHALHNSLSPAYGGAEFAELPSQAVPTDAHHPTPHHQQPAYPGPKEYLLP     1370
Rhesus_ENSMMUT00000028918.4     SSGSFEKKPDLHALHNSLSPAYGGAEFAELPSQAVPTDAHHPTPHHQQPAYPGPKEYLLP     1209
Rhesus_ENSMMUT00000028916.3     SSGSFEKKPDLHALHNSLSPAYGGAEFAELPSQAVPTDAHHPTPHHQQPAYPGPKEYLLP     1370
Rhesus_ENSMMUT00000075204.2     SSGSFEKKPDLHALHNSLSPAYGGAEFAELPSQAVPTDAHHPTPHHQQPAYPGPKEYLLP     1351
                                *.****************.************.*** ** *** *****************

Mouse_ENSMUST00000186548.7      KVPQLHPASRDPSPFAQSSSCYNRSIKQEPIDPLTQAESIPRDSAKMSRTPLPEASQNGG     1438
Human_ENST00000409262.8         KAPLLHSVSRDPSPFAQSSNCYNRSIKQEPVDPLTQAEPVPRDAGKMGKTPLSEVSQNGG     1430
Rhesus_ENSMMUT00000028918.4     KAPPIHSVSRDPSPFAQSSNCYNRSIKQEPVDPLTQAEPVPRDAGKMGKTPLPEVSQNGG     1269
Rhesus_ENSMMUT00000028916.3     KAPPIHSVSRDPSPFAQSSNCYNRSIKQEPVDPLTQAEPVPRDAGKMGKTPLPEVSQNGG     1430
Rhesus_ENSMMUT00000075204.2     KAPPIHSVSRDPSPFAQSSNCYNRSIKQEPVDPLTQAEPVPRDAGKMGKTPLPEVSQNGG     1411
                                *.* :* .***********.**********:******* :***:.**.:*** *.*****

Mouse_ENSMUST00000186548.7      PSHLWGQYSGGPSMSPKRTNSVGGNWGVFPPGESPTIVPDKLNSFGASCLTPSHFPESQW     1498
Human_ENST00000409262.8         PSHLWGQYSGGPSMSPKRTNGVGGSWGVFSSGESPAIVPDKLSSFGASCLAPSHFTDGQW     1490
Rhesus_ENSMMUT00000028918.4     PSHLWGQYSGGPSMSPKRTNSVGGSWGVFSSGESPALIPDKLSSFGASCLAPSHFTDGQW     1329
Rhesus_ENSMMUT00000028916.3     PSHLWGQYSGGPSMSPKRTNSVGGSWGVFSSGESPALIPDKLSSFGASCLAPSHFTDGQW     1490
Rhesus_ENSMMUT00000075204.2     PSHLWGQYSGGPSMSPKRTNSVGGSWGVFSSGESPALIPDKLSSFGASCLAPSHFTDGQW     1471
                                ********************.***.****  ****:::****.*******:**** :.**

Mouse_ENSMUST00000186548.7      GLFTGEGQQSAPHAGARLRGKPWSPCKFGNGTSALTGPSLTEKPWGMGTGDFNPALKGGP     1558
Human_ENST00000409262.8         GLFPGEGQQAASHSGGRLRGKPWSPCKFGNSTSALAGPSLTEKPWALGAGDFNSALKGSP     1550
Rhesus_ENSMMUT00000028918.4     GLFPGEGQQAASHSGGRLRGKPWSPCKFGNSTSTLTGPGLTEKPWGLGAGDFNSALKGGP     1389
Rhesus_ENSMMUT00000028916.3     GLFPGEGQQAASHSGGRLRGKPWSPCKFGNSTSTLTGPGLTEKPWGLGAGDFNSALKGGP     1550
Rhesus_ENSMMUT00000075204.2     GLFPGEGQQAASHSGGRLRGKPWSPCKFGNSTSTLTGPGLTEKPWGLGAGDFNSALKGGP     1531
                                *** *****:* *:*.**************.**:*:**.******.:*:**** ****.*

Mouse_ENSMUST00000186548.7      GFQDKLWNPVKVEEGRIPTPGANPLDKAWQAFGMPLSSNEKLFGALKSEEKLWDPFSLEE     1618
Human_ENST00000409262.8         GFQDKLWNPMKGEEGRIPAAGASQLDRAWQSFGLPLGSSEKLFGALKSEEKLWDPFSLEE     1610
Rhesus_ENSMMUT00000028918.4     GFQDKLWNPMKGEEGRIPAAGASQLDKAWQSFGLSLGSSEKLFGALKSEEKLWDPFSLEE     1449
Rhesus_ENSMMUT00000028916.3     GFQDKLWNPMKGEEGRIPAAGASQLDKAWQSFGLSLGSSEKLFGALKSEEKLWDPFSLEE     1610
Rhesus_ENSMMUT00000075204.2     GFQDKLWNPMKGEEGRIPAAGASQLDKAWQSFGLSLGSSEKLFGALKSEEKLWDPFSLEE     1591
                                *********:* ******:  **. **:***:**: *.*.********************

Mouse_ENSMUST00000186548.7      GTAEEPPSKGVVKEEKSGPTVEEDEEELWSDSEHNFLDENIGGVAVAPAHCSILIECARR     1678
Human_ENST00000409262.8         GPAEEPPSKGAVKEEKGGGGAEEEEEEELWSDSEHNFLDENIGGVAVAPAHGSILIECARR     1670
Rhesus_ENSMMUT00000028918.4     GPAEEPPSKGAVKEEKGGGGAEEEEEEELWSDSEHNFLDENIGGVAVAPAHGSILIECARR     1509
Rhesus_ENSMMUT00000028916.3     GPAEEPPSKGAVKEEKGGGGAEEEEEEELWSDSEHNFLDENIGGVAVAPAHGSILIECARR     1670
Rhesus_ENSMMUT00000075204.2     GPAEEPPSKGAVKEEKGGGGAEEEEEEELWSDSEHNFLDENIGGVAVAPAHGSILIECARR     1651
                                * ********.*****.*  .**:************************** *********

Mouse_ENSMUST00000186548.7      ELHATTPLKKPNRCHPTRISLVFYQHKNLNQPNHGLALWEAKMKQLAERARQRQEEAARL     1738
Human_ENST00000409262.8         ELHATTPLKKPNRCHPTRISLVFYQHKNLNQPNHGLALWEAKMKQLAERARARQEEAARL     1730
Rhesus_ENSMMUT00000028918.4     ELHATTPLKKPNRCHPTRISLVFYQHKNLNQPNHGLALWEAKMKQLAERARARQEEAARL     1569
Rhesus_ENSMMUT00000028916.3     ELHATTPLKKPNRCHPTRISLVFYQHKNLNQPNHGLALWEAKMKQLAERARARQEEAARL     1730
Rhesus_ENSMMUT00000075204.2     ELHATTPLKKPNRCHPTRISLVFYQHKNLNQPNHGLALWEAKMKQLAERARARQEEAARL     1711
                                *************************************************** ********

Mouse_ENSMUST00000186548.7      GLGQQEAKLYGKKRKWGGAMVAEPQHKEKKGAIPTRQALAMPTDSAVTVSSYAYTKVTGP     1798
Human_ENST00000409262.8         GLGQQEAKLYGKKRKWGGTVVAEPQQKEKKGVVPTRQALAVPTDSAVTVSSYAYTKVTGP     1790
Rhesus_ENSMMUT00000028918.4     GLGQQEAKLYGKKRKWGGAMVAEPQQKEKKGVVPTRQALAVPTDSAVTVSSYAYTKVTGP     1629
Rhesus_ENSMMUT00000028916.3     GLGQQEAKLYGKKRKWGGAMVAEPQQKEKKGVVPTRQALAVPTDSAVTVSSYAYTKVTGP     1790
Rhesus_ENSMMUT00000075204.2     GLGQQEAKLYGKKRKWGGAMVAEPQQKEKKGVVPTRQALAVPTDSAVTVSSYAYTKVTGP     1771
                                ******************::*****:*****.:*******:*******************

Mouse_ENSMUST00000186548.7      YSRWI*      1803
Human_ENST00000409262.8         YSRWI*      1795
Rhesus_ENSMMUT00000028918.4     YSRWI*      1634
Rhesus_ENSMMUT00000028916.3     YSRWI*      1795
Rhesus_ENSMMUT00000075204.2     YSRWI*      1776
                                ******
```

CYS rich region

Catalytic residue

Large low complexity insert
